# Modifiers of genetic dominance at the Arabidopsis self-incompatibility locus retain proto-miRNA features and act through non-canonical gene silencing pathways

**DOI:** 10.1101/2024.05.01.591913

**Authors:** Rita A. Batista, Eléonore Durand, Monika Mörchen, Jacinthe Azevedo-Favory, Samson Simon, Manu Dubin, Vinod Kumar, Eléanore Lacoste, Corinne Cruaud, Christelle Blassiau, Matteo Barois, Anne-Catherine Holl, Chloé Ponitzki, Nathalie Faure, William Marande, Sonia Vautrin, Isabelle Fobis-Loisy, Jean-Marc Aury, Sylvain Legrand, Ute Krämer, Thierry Lagrange, Xavier Vekemans, Vincent Castric

**Affiliations:** Univ. Lille, CNRS, UMR 8198 – Evo-Eco-Paleo, F-59000 Lille, France; CNRS, Laboratoire Génome et Développement des Plantes, UMR 5096, 66860 Perpignan, France, and Université Perpignan Via Domitia, Laboratoire Génome et Développement des Plantes, UMR 5096, F-66860 Perpignan, France; Department of Molecular Genetics and Physiology of Plants, Faculty of Biology and Biotechnology, Ruhr University Bochum, D-44801 Bochum, Germany; Génomique Métabolique, Genoscope, Institut François Jacob, CEA, CNRS, Univ Evry, Université Paris-Saclay, 91057 Evry, France; Genoscope, Institut François Jacob, CEA, Université Paris-Saclay, Evry, 91057, France; INRAE - CNRGV French Plant Genomic Resource Center - Castanet Tolosan, France; Laboratoire Reproduction et Développement des Plantes, Université de Lyon, ENS de Lyon, UCB Lyon 1, CNRS, INRAE, F-69342 Lyon, France

## Abstract

Self-incompatibility in flowering plants is a common mechanism that prevents self-fertilization and promotes outcrossing. In Brassicaceae, the self-incompatibility locus is highly diverse, with many alleles arranged in a complex dominance hierarchy and exhibiting monoallelic expression in heterozygote individuals. Monoallelic expression of the pollen self-incompatibility gene is achieved through the action of sRNA precursors that resemble miRNAs, although the underlying molecular mechanisms remain elusive. Here, we engineered *Arabidopsis thaliana* lines expressing components of the *Arabidopsis halleri* self-incompatibility system, and used a reverse genetics approach to pinpoint the pathways underlying the function of these sRNA precursors. We showed that they trigger a robust decrease in transcript abundance of the recessive self-incompatibility genes, but not through the canonical transcriptional or post-transcriptional gene silencing pathways. Furthermore, we observed that single sRNA precursors are typically processed into hundreds of sRNA molecules with a variety of sizes, abundance levels and ARGONAUTE loading preferences. Our results suggest that these seemingly arbitrary processing characteristics are essential for establishing the self-incompatibility dominance hierarchy, as they enable a single sRNA precursor from a dominant allele to effectively repress multiple recessive alleles, thus providing a unique example of how small RNAs mediate gene silencing within a highly complex regulatory network.

**Graphical abstract:** 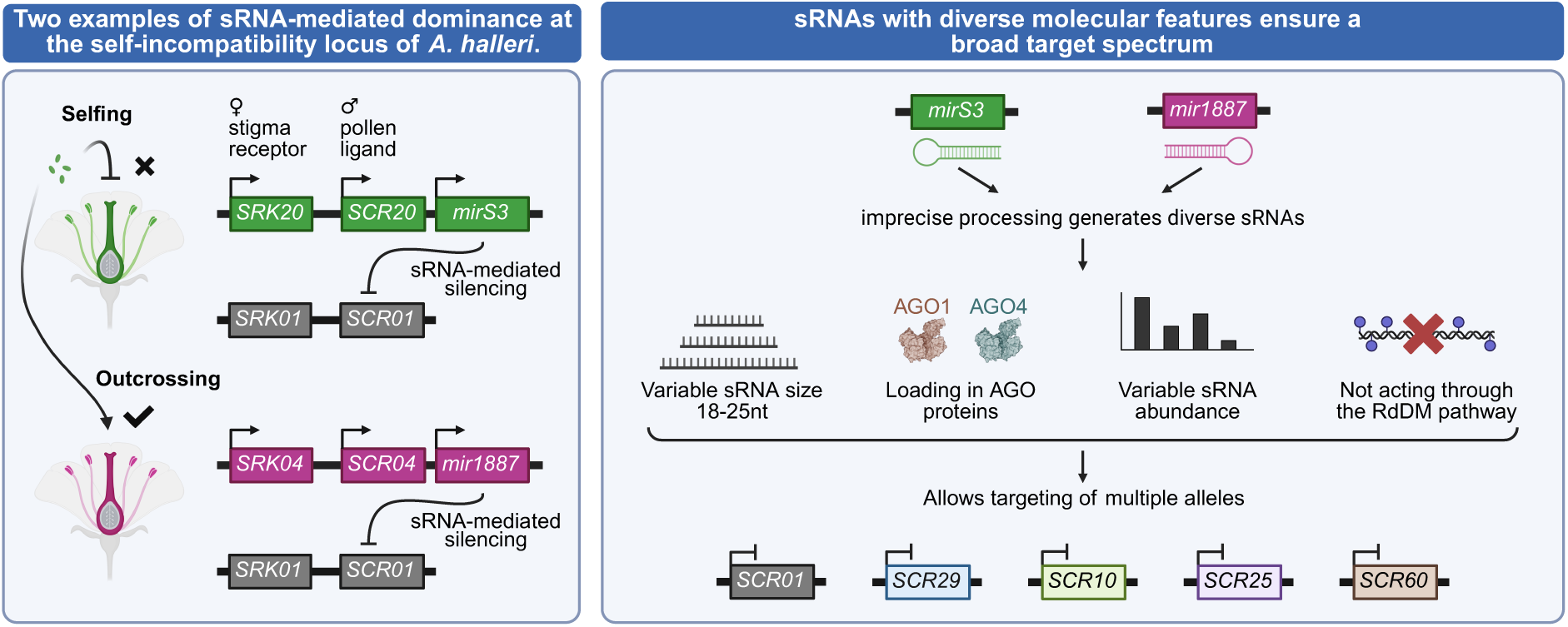

## Introduction

Self-incompatibility is a widespread self-recognition mechanism in hermaphroditic flowering plants that prevents selfing and enforces outcrossing. In Brassicaceae, self-pollen recognition is controlled at the sporophytic level by a single non-recombining self-incompatibility locus (*S-*locus) containing two protein-coding genes: the female component encoding a trans-membrane receptor (SRK) expressed in papillae cells, and the male component consisting of an SCR peptide (referred to as SP11 in the Brassica nomenclature) produced in the anther tapetum and deposited on the surface of the pollen grains. The SRK and SCR proteins function as a receptor-ligand system, such that pollen germination is stopped when a pair of cognate SRK and SCR proteins interact upon pollen deposition on the pistil, ultimately preventing self-fertilization (1–5). The *S-*locus has been subject to an intense long-term balancing selection, which favors the maintenance of a high number of distinct *S-*alleles (6–9). Although heterozygosity is typically very high at the *S-*locus, monoallelic expression of the male *SCR* gene is the rule, following a transitive and mostly linear dominance hierarchy among *S-*alleles (10–12). These dominance interactions maximize reproductive success by ensuring that in a diploid individual only one of the two *SCR* alleles is expressed, thus preventing pollen rejection by females that share the same allelic specificity as the repressed, recessive *SCR*, and thereby increasing the number of compatible female partners (13). In *Arabidopsis lyrata* and *A. halleri*, respectively, 56 and 43 *S-*alleles have been documented so far (7, 14), which are divided into four main dominance classes - class IV being the most dominant and class I the most recessive (15, 16).

Underlying these dominance interactions are dominance modifiers - genetic elements dedicated to altering dominance relationships between other alleles in *trans* (17). In the particular context of the self-incompatibility system these dominance modifiers are located in the *S*-locus and take the form of sRNA-generating loci that are hypothesized to modulate the expression of the *SCR* gene_(18). Similarly to miRNAs genes, these loci are transcribed into an RNA molecule that is predicted to fold and form a hairpin structure, which is subsequently cleaved into sRNAs. In *A. halleri*, eight distinct families of sRNA precursors, defined based on sequence similarity, are predicted to regulate dominance interactions both between and within the four *S*-allele dominance classes (19). Interestingly, in Brassica where only two dominance classes exist, only two sRNA precursor families have been identified (12, 20, 21), suggesting that there is an association between the complexity of the dominance network and the total number of dominance modifiers. The two sRNA precursors of Brassica (*Smi* and *Smi2*) have each been suggested to produce a 24-nt sRNA with sequence similarity to the 5’ region of *SCR.* The presence of this sRNA is associated with decreased expression of the recessive *SCR* allele, as well as deposition of DNA methylation within and around the targeted region in tapetum cells (12, 20, 21). This is reminiscent of what is observed in several plant species where 21/24-nt sRNAs derived from miRNAs genes or inverted repeats can be loaded onto proteins of the ARGONAUTE4 (AGO4) clade and subsequently elicit *de novo* DNA methylation at target loci, such as transposable elements or protein-coding genes, through the RNA-directed DNA Methylation (RdDM) pathway (22–26). Similar mechanisms could be utilized by *Smi* and *Smi2*; nevertheless, this hypothesis was never empirically tested.

In Brassica, the dominance hierarchy of *S*-alleles was proposed to rely on the combinatorial effect of single nucleotide variants at the *Smi* and *Smi2* sequences and at their respective *SCR* target sites (21). In Arabidopsis, where more dominance classes exist, the dominance network is hypothesized to rely on two properties: i) recessive alleles carry more target sites and these sites are more generalist (*i.e.* they are targeted by more sRNA precursors families than dominant alleles); and ii) sRNA precursors of dominant alleles are more generalist (*i.e.* they target a higher number of alleles than those sRNA derived from precursors in more recessive alleles) (19). Despite this observation, the molecular features allowing sRNA precursors to target multiple alleles and have a broad targeting spectrum remain to be identified. In addition, among the eight families of sRNA precursors previously predicted to control the dominance hierarchy in Arabidopsis, only one has been subject to formal experimental validation (19).

In this study, we aimed at identifying the molecular pathway underlying the sRNA-mediated dominance interactions between *S-*alleles in Arabidopsis. To achieve this, we engineered and characterized a series of *A. thaliana* lines recapitulating the self-incompatibility phenotype of *A. halleri*, and used these lines to show that two different sRNA precursors from dominant *A. halleri S*-loci trigger a decrease in abundance of recessive *SCR* transcripts. We thus provide direct proof that this heterologous approach can be used to validate sRNA precursor function, both at the phenotypic and at the molecular level. This system also allowed us to investigate if key components of the canonical RdDM pathway, such as the ARGONAUTE proteins AGO4 and AGO6, and subunits of the DNA-dependent RNA polymerases POLIV and POLV, are required for the repression of *SCR* transcript by *S-*locus sRNA precursors, and our analyses demonstrate that the two selected sRNA precursors do not act through the canonical RdDM pathway.

Additionally, we reveal that despite their structural similarity to miRNAs, *S-*locus-derived sRNA precursors produce a large population of sRNAs with different sizes, sequences, ARGONAUTE loading preferences and target sites. We show that this molecular heterogeneity is important to maximize the number of targeted *S*-alleles, thus allowing one dominant allele to silence multiple recessive alleles at the same time. Overall, our results show how the distinctive molecular features of *S*-locus sRNAs contribute to effective gene silencing within a complex regulatory network, shedding light on their role in shaping dominance interactions of an important reproductive phenotype.

## Results

### The sRNA precursors Ah04mir1887 and Ah20mirS3 control dominance interactions at the S-locus by decreasing the transcript level of the recessive SCR01 allele

To study in detail the molecular mechanisms controlling the activity of dominance modifiers in the self-incompatibility system of *A. halleri*, we first recreated the self-incompatibility reaction of *A. halleri* in the selfing species *A. thaliana*, since the latter is a more tractable study model. To do this, we expressed the *SRK01* receptor and its cognate *SCR01* ligand in female and male parents, respectively (19, 27, 28). Manual crosses between these plants revealed that the germination of pollen from a *SCR01*-expressing male was abolished when deposited on pistils of an *SRK01*-expressing female, the hallmark of a self-incompatible reaction **(Fig. 1C and D, WT panels)**. Given the challenges faced in prior studies to reconstruct this reaction in the Col-0 ecotype, all *A. thaliana* plants used in this study are of the C24 ecotype, where the self-incompatibility can be faithfully recreated as shown both in this, and in other studies (27, 29, 30).

**Figure 1.**
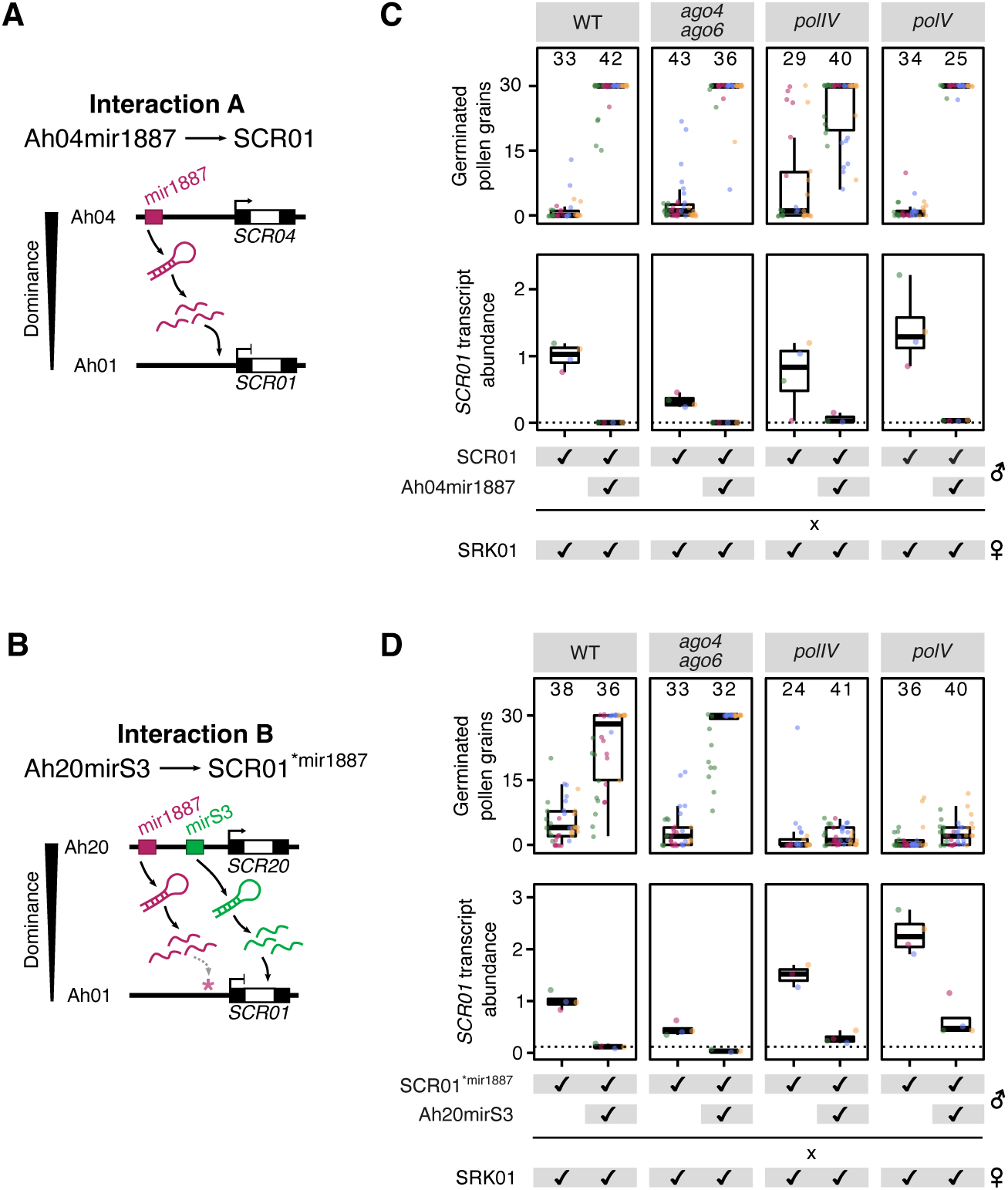
The RdDM pathway is not required for the activity of Ah04mir1887 and Ah20mirS3. **(A,B)** Dominance interactions studied in this paper. The black and white regions within *SCR* genes correspond to exons and introns, respectively. **(A)** Interaction A focuses on the role of the sRNA precursor Ah04mir1887 in mediating the dominance of Ah04 over Ah01. **(B)** Interaction B focuses on the role of Ah20mirS3 in mediating the dominance of Ah20 over Ah01. *S-*locus sRNA precursors consist of inverted repeats that form hairpins which are processed into multiple sRNAs showing sequence homology to specific *SCR* regions: Ah04mir1887 sRNAs show homology to the 5’ region region of *SCR01*, while Ah20mirS3 sRNAs show homology to the intronic region (white) of *SCR01*. Because sRNAs produced from the Ah20mir1887 precursor show homology to the wt *SCR01* 5’ region region, and to study the specific effect of Ah20mirS3 in Interaction B, the *SCR01^*mir1887^* line containing five point mutations in the *SCR01* 5’ region was created, which disrupts the homology with Ah20mir1887 sRNAs. **(C,D)** Phenotypic and expression assay evaluating the role of key RdDM components on the action of Ah04mir1887 **(C)**, and Ah20mirS3 **(D)**. Crosses between *SRK01*-expressing mothers and *SCR01*- expressing fathers were performed in wt and RdDM mutant backgrounds in the absence or presence of the respective sRNA precursor. The number of germinated pollen grains of the indicated genotype was counted on pistils of SRK01 mothers. *SCR01* expression in immature buds of the indicated genotype was measured by RT-qPCR and normalized to the wt level. Five biological replicates (represented by the differently colored dots) were analyzed for pollen germination and *SCR01* expression. Numbers on top of columns indicate the total number of siliques sampled for pollen germination assays.

Among the eight known families of *A. halleri* sRNA precursor loci, only one - mirS3 - has been experimentally validated to regulate dominance between two distinct *S*-alleles: specifically, the dominance of allele Ah20 over Ah01, mediated by Ah20mirS3 sRNAs, which target the *SCR01* intronic region **(Fig. 1B)** (19). In this study, we set out to further investigate the molecular properties of Ah20mirS3, as well as examine if the yet uncharacterized mir1887 family could also function as a dominance modifier. For this, we focused on testing the role of the sRNA precursor Ah04mir1887 in controlling the dominance of allele Ah04 over Ah01, whose sRNAs are predicted to target the promoter region of *SCR01*, immediately upstream of the TSS **(Fig. 1A**, **Fig. 1 S1**). We thus generated transgenic lines expressing Ah04mir1887, introgressed this transgene into an *SCR01*-expressing male, and used these plants to pollinate *SRK01*-expressing females, testing whether the presence of the Ah04mir1887 sRNA precursor locus could transform the previously incompatible interaction between *SCR01* and *SRK01* into a compatible one **(Fig. 1C)**. In the particular case of Ah20mirS3, the transgenic line previously generated by Durand et al. (19) and used here expresses both Ah20mirS3 and Ah20mir1887 because of their close physical proximity within the Ah20 allele **(Fig. 1B)**. To specifically isolate the impact of Ah20mirS3 in the dominance interaction of Ah20 over Ah01, we developed the *SCR01^*mir^*^1887^ transgenic line, which contains five point mutations designed to fully disrupt the slight homology observed between some Ah20mir1887 sRNAs and *SCR01* **(Fig. 1B**; **Fig. 1 S2)**. We then used this modified *SCR01^*mir^*^1887^ line for all subsequent tests on the functional role of Ah20mirS3.

We observed that combining either Ah04mir1887 (Interaction A), or Ah20mirS3 (Interaction B) with *SCR01* results in a switch from an incompatible to a compatible pollen germination phenotype, which is accompanied by a dramatic decrease in the abundance of the *SCR01* transcript, as measured by RT-qPCR **(Fig. 1C and D**, **WT panels)**. This shows that both Ah04mir1887 and Ah20mirS3 act as dominance modifiers in their respective allelic interactions, and that they accomplish this by decreasing *SCR01* transcript abundance, thus preventing SRK-SCR cognate recognition and allowing pollen germination to occur. Moreover, these results demonstrate that *SCR* transcript abundance can be modulated by multiple sRNA precursor families that use distinct target sites in the *SCR* locus.

### Decrease in SCR01 transcript abundance does not depend on RdDM

Next, we aimed at elucidating the exact molecular mechanism responsible for the reduction of *SCR01* transcript abundance triggered by the two *S*-locus sRNA precursors. Given their resemblance to miRNAs genes, one potential scenario is that sRNAs produced by Ah04mir1887 and Ah20mirS3 could trigger post-transcriptional gene silencing (PTGS) of the *SCR01* transcript. However, since these sRNAs are predicted to target the promoter and intronic region of *SCR01*, and since the substrate of PTGS is generally thought to be the mature mRNA (31), this does not support PTGS as the pathway responsible for the decrease in *SCR01* transcript abundance. An alternative scenario is that Ah04mir1887 and Ah20mirS3 sRNAs could trigger the RNA-directed DNA methylation pathway (RdDM), resulting in the deposition of DNA methylation in their respective *SCR01* target sites, and reduced *SCR01* transcription. Interestingly, previous studies have established that the silencing of recessive alleles in the homologous self-incompatibility system of *Brassica* relies on sRNA-mediated DNA methylation, although the specific pathway underlying this process remains elusive (12, 20, 21). Thus, we focused our attention on the RdDM pathway, and set out to test if mutating key components of this pathway would prevent Ah04mir1887 and Ah20mirS3 from reducing *SCR01* transcript abundance. To circumvent the absence of RdDM mutant alleles in the C24 ecotype, we generated and validated C24 *ago4 ago6* CRISPR/Cas9 mutant alleles **(Fig. 1 S3)**, and used the previously published C24 *polIV* and *polV* knock-out alleles (32).

To our surprise, none of the RdDM mutants impaired the activity of the tested sRNA precursors. In the case of Ah04mir1887 (Interaction A), *SCR01* transcript abundance was reduced in all mutant backgrounds to a similar level as in wild type, which was reflected by a compatible pollen germination phenotype **(Fig. 1C)**. Similarly, the presence of *Ah20mirS3* (Interaction B) was associated with a decrease in *SCR01* transcript in all mutant backgrounds, showing that this sRNA precursor is still functional when RdDM is impaired **(Fig. 1D)**. And while in the *ago4 ago6* mutant background the reduction in *SCR01* transcript led to a compatible pollen phenotype, in both *polIV* and *polV* mutant backgrounds the reduction in *SCR01* transcript level was sufficient to elicit a compatible pollen phenotype **(Fig. 1D)**. This is likely explained by the fact that in Interaction B the basal *SCR01* transcript level is increased in both *polIV* and *polV* mutant backgrounds when compared with wild type **(Fig. 1D**, ***polIV* and *polV* panels vs. WT panel)**, pointing to an effect of these mutants on the expression of the *SCR01* transgene. This suggests that the *SCR01^*mir^*^1887^ construct might be intrinsically targeted for repression by RdDM, a phenomenon often observed in transgenic lines (33) (note that the *SCR01* and *SCR01^*mir^*^1887^ transgenes used in Interaction A and B, respectively, correspond to two independent transgenic lines with different insertion sites; **see Methods section**). Indeed, our data shows that a strong reduction of *SCR01* transcript abundance is required for a compatible pollen germination phenotype, since even very low levels of transcript can trigger an incompatibility reaction **(Fig. 1 S4)**. Together, these results indicate that neither PTGS nor RdDM, at least not in their canonical forms, appear to be responsible for the reduction in *SCR01* transcript abundance triggered by *S*-locus sRNA precursors. Despite this, both Ah04mir1887 and Ah20mirS3 strongly and robustly decrease *SCR01* transcript levels, which is necessary to abolish the self-incompatibility reaction, as shown here.

### Ah04mir1887 and Ah20mirS3 hairpins undergo heterogeneous processing to generate diverse sRNA populations

*S*-locus sRNA precursors, including those of the mir1887 and mirS3 families studied here, consist of inverted repeats that are transcribed and predicted to form a RNA hairpin structure containing a 5’ arm, a terminal loop and a complementary 3’ arm **(Fig. 2A)** (19), akin to the structures formed by precursor miRNA (pri-miRNA) molecules. To delve deeper into the molecular properties of Ah04mir1887 and Ah20mirS3, and to explore potential non-canonical pathways through which these precursors may act, we conducted several sRNA sequencing experiments on both transgenic *A. thaliana* and wild type *A. halleri* plants harboring Ah04mir1887 and Ah20mirS3. The resulting data allowed us to compile a comprehensive database encompassing all sRNAs molecules sequenced for each precursor, irrespective of their species origin. Using these data, we observed that dicing of *S-* locus sRNA precursor hairpins leads to numerous sRNAs – at least 106 in the case of Ah04mir867, and 224 in the case of Ah20mirS3 **(Fig. 2B)**. These sRNAs range from 18 to 25 nt in length, with no clear predominance of a specific length **(Fig. 2 S1 A-B)**, and their abundances vary by several orders of magnitude **(Fig. 2B**, **Fig. 2 S1 C-D)**.

**Figure 2.**
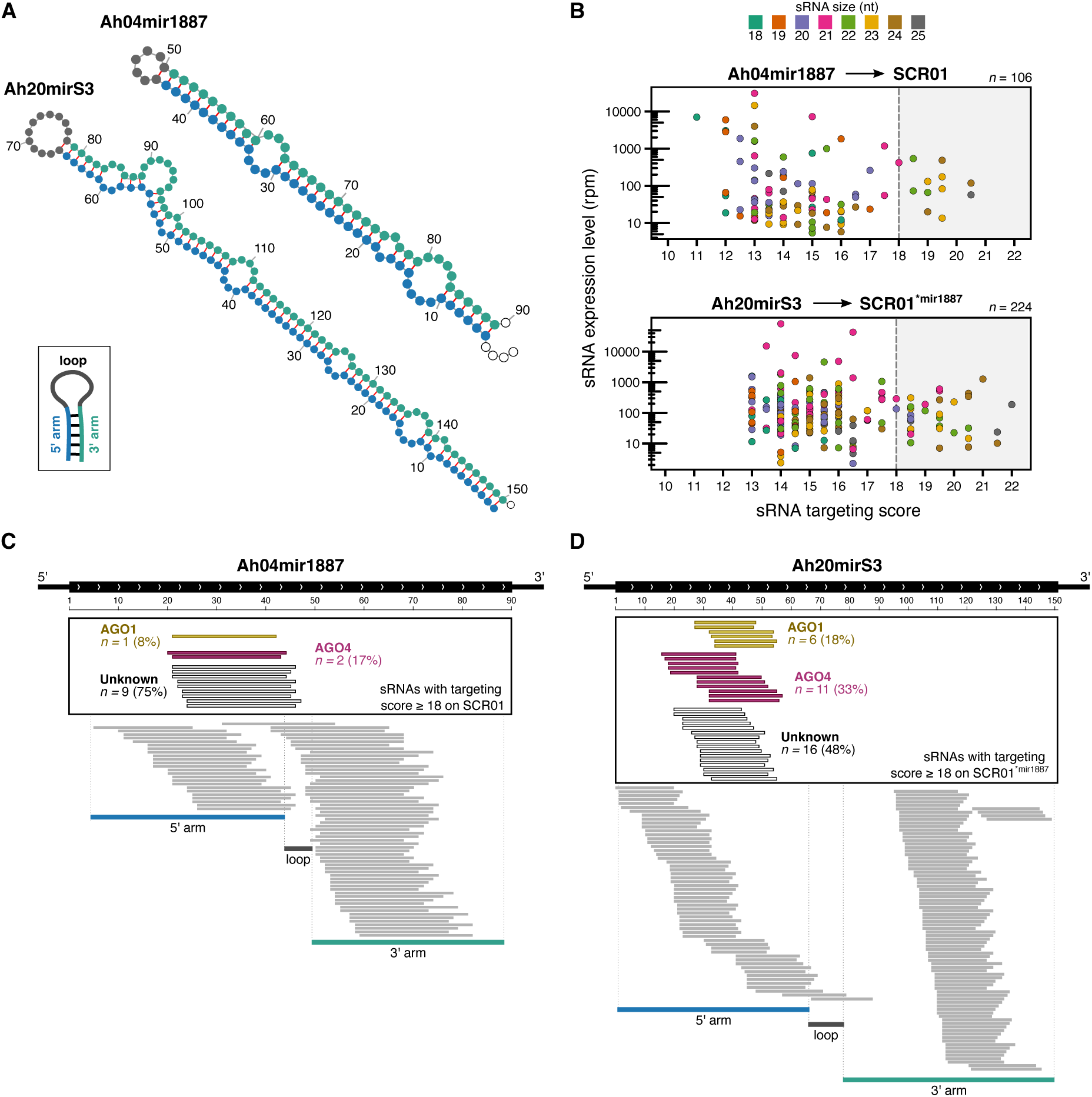
Ah04mir1887 and Ah20mirS3 hairpins are processed into numerous sRNAs. **(A)** Predicted RNA hairpin structures of the *S-*locus sRNA precursors Ah04mir1887 and Ah20mirS3. **(B)** Expression of Ah04mir1887 and Ah20mirS3 sRNAs as a function of their targeting score against *SCR01.* The dot color corresponds to the size of the sRNA. The grey box highlights sRNAs predicted to induce a reduction in *SCR01* transcript abundance (score ≥ 18). *n* corresponds to the total number of unique sRNAs identified for each sRNA precursor. **(C-D)** Genomic representation of the Ah04mir1887 and Ah20mirS3 loci, and their respective sRNAs. The black rectangle highlights sRNAs predicted to induce a reduction in *SCR01* transcript abundance (score ≥ 18). AGO loading information for each functional sRNA is indicated by the sRNA color (yellow – sRNA loaded predominantly in AGO1; magenta – sRNA loaded predominantly in AGO4; white – sRNAs with unknown loading profile), **see Methods section *ARGONAUTE immunoprecipitation and sRNA sequencing*** for more details.

Given the large population of sRNAs produced by a single precursor, we aimed at evaluating the functional relevance of each sRNA in controlling the abundance of the *SCR01* transcript. For this, we used a previously published algorithm (19), which attributes a targeting score based on the homology between a single sRNA and the *SCR* genomic region. A previous study has demonstrated that in multiple *A. halleri S*-locus heterozygotes, a targeting score equal to, or above 18 is associated with phenotypic dominance and a sharp decrease of the recessive *SCR* transcript level (10). Using this algorithm, we found that 12 out of the total 106 sRNAs produced by Ah04mir1887, and 33 out of the total 224 sRNAs produced by Ah20mirS3, have a base-pairing score of ≥ 18 and are thus predicted to target *SCR01*. This subset of putatively functional sRNAs displays a variety of sizes and expression levels **(Fig. 2B**, **see grey box)**.

To further assess whether these molecular features are reproducible between the native *A. halleri* context and the transgenic *A. thaliana* context, we performed the same analyses as above, but using only the sRNAs detected in *A. halleri* samples **(Fig. 2 S2)**. Although fewer sRNAs were identified in *A. halleri*, their size distribution, abundances, and targeting spectrum closely mirror those observed in the combined sRNA dataset from both species **(Fig. 2 S2 A-F)**. We hypothesize that the reduced number of sRNAs detected in *A. halleri* is primarily due to the lower sequencing effort in this species compared to *A. thaliana* **(Fig. 2 S2 G)**, suggesting that the total number of sRNAs identified for each precursor is highly dependent on sequencing depth, and that the saturation point was not reached in *A. halleri*. Overall, these data indicate that the transgenic *A. thaliana* system accurately recapitulates the features observed in the native *A. halleri* context, further supporting the use of *A. thaliana* as a more tractable system to study the molecular bases of self-incompatibility reactions. Based on these findings, we decided to proceed with the initially compiled sRNA database, which includes the sum of sRNAs sequenced in both *A. halleri* and *A. thaliana*, for further downstream analyses in this manuscript.

To better understand the AGO loading preferences for each sRNA we performed AGO immunoprecipitation (IP) assays, followed by sRNA sequencing in the Ah04mir1887 and Ah20mirS3 *A. thaliana* transgenic lines. This showed that functional sRNAs (base-pairing score of ≥ 18) produced from either precursor are loaded both into AGO1 (characteristic of the PTGS pathway) and AGO4 (characteristic of the RdDM pathway) **(Fig. 2 C-D)**. Notwithstanding this, a large part of these functional sRNAs (75% in Ah04mir1887 and 48% in Ah20mirS3) are not found to be loaded in either of these two AGO proteins, consistent with our observations that both the canonical PTGS and RdDM pathways are unlikely to underlie the activity of these sRNA precursors. We also noted that the majority of sRNAs loaded in AGO1 have a 5’ terminal uracil, reflecting the known AGO1 preference for binding these sRNAs (34). However, sRNAs loaded into AGO4, or with an unknown loading pattern, do not show a significant bias towards a specific 5’ nucleotide **(Fig. 2 S3)**.

Despite the structural similarity between *S*-locus sRNA precursors and miRNAs, and the existence of shared features, such as the presence of mature 21-nt sRNAs and loading of some sRNAs into AGO1, it is evident that *S*-locus sRNA precursors deviate from the typical characteristics associated with canonical miRNAs, and cannot be characterized as such. Instead, our data show that Ah04mir1887 and Ah20mirS3 have an unusually staggered hairpin processing pattern with numerous sRNAs derived from both the 5’ and 3’ arms, which differ in size, abundance level and ability to associate with AGO proteins.

### Heterogeneous processing of S-locus sRNA precursors allows targeting of multiple S-alleles

We hypothesized that this heterogeneity in hairpin processing pattern could play an important role in the context of the complex *S-*locus dominance hierarchy by allowing a single sRNA precursor to exert dominance over multiple other alleles. To test this, we expanded our analysis of the function of Ah04mir1887 and Ah20mirS3 beyond the interaction with allele Ah01, and looked at their role within the *A. halleri* dominance network **(Fig. 3)**.

**Figure 3.**
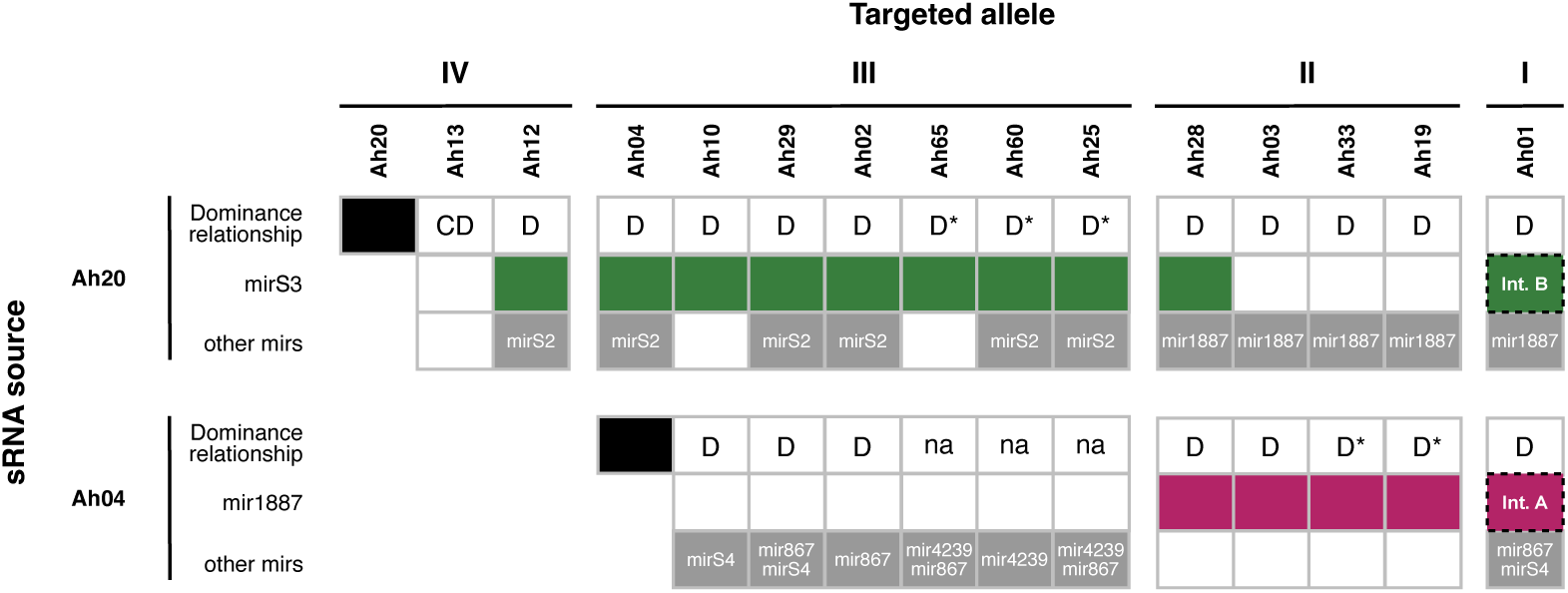
Ah04mir1887 and Ah20mirS3 underlie numerous *S-*allele dominance interactions. Schematic representation of the dominance relationship between alleles Ah20, Ah04 and other *S-* alleles in the current *A. halleri* dominance network. The dominance phenotypes observed are represented by D (dominant, as determined by phenotypic assays), CD (codominant, as determined by phenotypic assays), and na (no phenotypic data available). D* refers to a predicted dominance relationship, based on the difference between the dominance class of the sRNA source and the targeted allele (7, 16). The sRNA precursors underlying each dominance relationship are highlighted in green (mirS3), magenta (mir1887), or grey (other mirs). The specific interactions studied earlier in this manuscript – Interaction A and Interaction B - are highlighted by a dashed box.

We compiled previously published phenotypic data obtained by controlled reciprocal crosses between different *A. halleri S-*locus heterozygotes to determine the position of allele Ah20 and Ah04 in the dominance hierarchy of a total of 15 *S-*alleles **(Fig. 3)** (10, 19, 35). From this, we inferred that Ah20 is dominant over 13 other *S-*alleles, with 10 of these interactions relying on the action of Ah20mirS3 **(Fig. 3)**, as determined by the sRNA-*SCR* targeting algorithm described above. Thus, Ah20mirS3 is not only able to mediate the repression of *SCR01*, but also the repression of at least nine other distinct *SCR* alleles. Similarly, Ah04mir1887 mediates the repression of at least five different *SCR* alleles that are recessive to Ah04 **(Fig. 3)**.

To test if the large population of sRNAs produced by these *S-*locus precursors is important in repressing multiple target alleles, we examined the role of each individual sRNA produced by both Ah04mir1887 and Ah20mirS3 **(Fig. 4)**: 11% of Ah04mir1887 sRNAs **(Fig. 4A**, **cluster 1)**, and 26% of Ah20mirS3 sRNAs **(Fig. 4B**, **cluster 1)** are predicted to target at least one *SCR* allele within the inferred dominance network presented in **Figure 3**. As is the case in Interaction A and B, there does not seem to be a clear correlation between abundance and sRNA functionality in any of the analyzed interactions **(Fig. 2B and Fig. 4**, **see cluster 1 vs. cluster 2)**. Nevertheless, sRNAs that are predicted to be functional have a larger size (median of 23 nt vs. 21 nt), and in the case of Ah20mirS3, this is also associated with a larger number of sRNAs being loaded in AGO proteins **(Fig. 4**, **see cluster 1 vs. cluster 2).** Additionally, according to our sRNA-*SCR* targeting algorithm, most *S-*alleles are redundantly targeted by multiple sRNAs **(Fig. 4)**.

**Figure 4.**
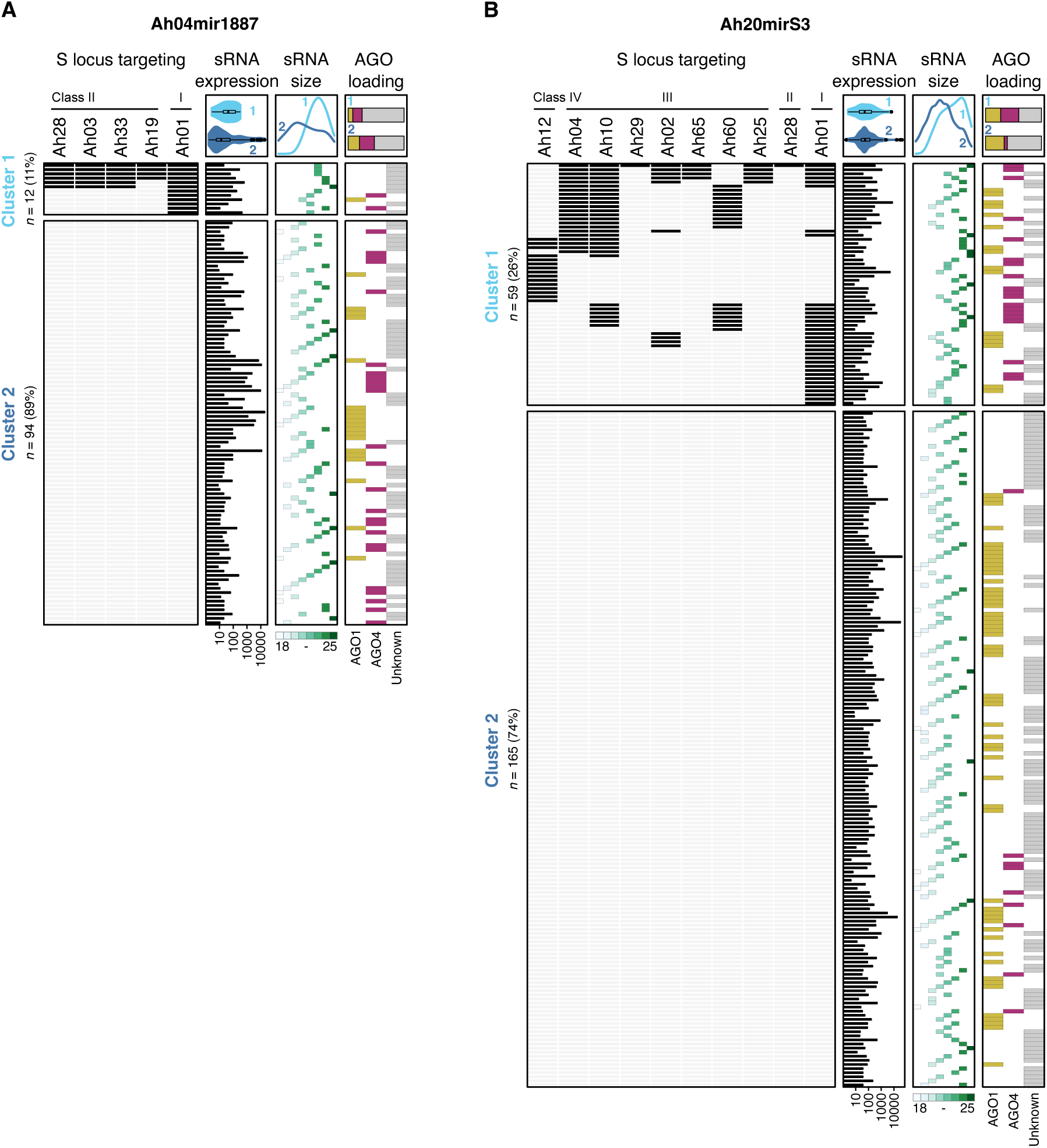
Ah04mir1887 and Ah20mirS3 produce multiple sRNAs predicted to target one or more *S-*alleles. Heatmap representing the *S-*loci targeted by Ah04mir1887-derived sRNAs **(A)** and Ah20mirS3-derived sRNAs **(B)**. A black rectangle indicates that a specific sRNA has a targeting score of ≥ 18 against the corresponding *S-*allele. Cluster 1 groups sRNAs that target at least one *S-*allele, while Cluster 2 groups sRNAs that have no targets in the represented *S-*alleles. sRNA expression levels, sRNA size and AGO loading patterns are indicated for each sRNA. Summary plots comparing these features in Cluster 1 and Cluster 2 are shown in the top panel. *n* corresponds to the number of sRNAs produced by each sRNA precursor.

When investigating in more detail the positional correspondence between sRNAs and the Ah04mir1887/Ah20mirS3 precursor hairpin structure we noticed that sRNAs are produced in similar amounts from both the 5’ and the 3’ arms of the hairpin **(Fig. 5)**. Each hairpin arm produces numerous sRNAs that show a staggered pattern, suggesting imprecise dicing and resulting in a population of sRNAs that show only a few nucleotide differences between them. In the case of Ah04mir1887, only the sRNAs produced from the 5’ hairpin arm are predicted to be functional against the *S-*alleles tested here **(Fig. 5A)**. Remarkably, in the case of Ah20mirS3, each hairpin arm shows functional specialization since sRNAs from the 5’ arm target all allele classes except Class IV, while sRNAs from the 3’ arm exclusively target Class IV and Class III alleles **(Fig. 5B)**. Together, these observations suggest that the high number and heterogeneous processing of sRNAs is a conserved and important feature of *S-* locus sRNA precursors which allows targeting of multiple alleles and thus underlies their generalism.

**Figure 5.**
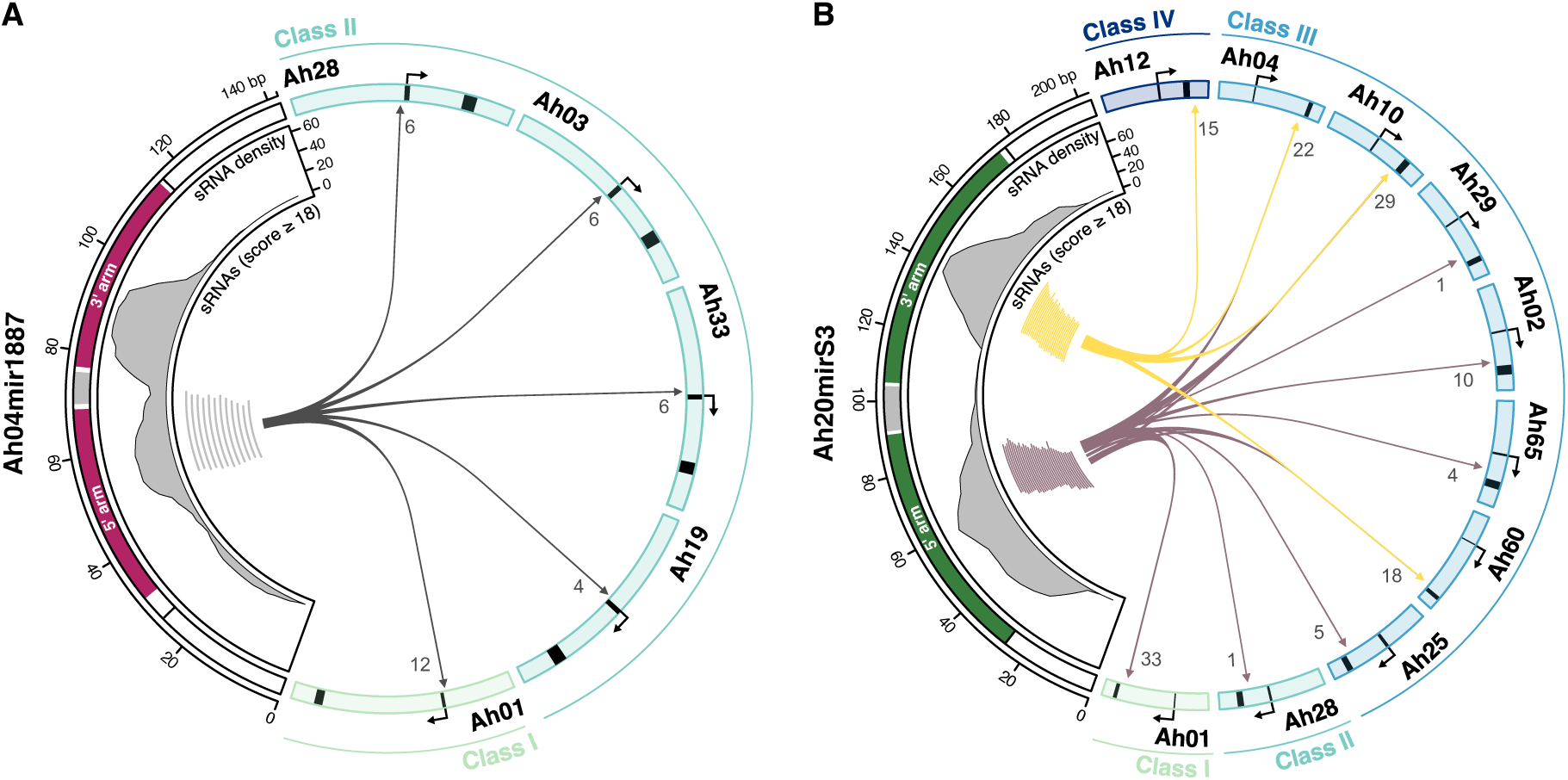
The targeting repertoire of *S-*locus sRNA precursors is enhanced by their heterogeneous processing pattern. Circos plot representing the positional relationship between sRNAs produced by Ah04mir1887 **(A)**, Ah20mirS3 **(B)** and the corresponding targeted *S-*alleles. The genomic regions harboring Ah04mir1887 and Ah20mirS3 are represented on the left side of the plots. The predicted 5’ and 3’ hairpin arms are represented in magenta (Ah04mir1887) and green (Ah20mirS3), and the sRNA density (number of sRNAs) is plotted along the sRNA precursor region. Individual sRNAs with a score of ≥ 18 are represented by the grey, yellow or dark purple segments. Arrowed segments represent the sRNA source - target relationship. The numbers at the end of these arrowed segments indicate the total number of sRNAs targeting that respective allele. The *SCR* genomic region of targeted *S-*alleles is represented on the right side and includes 1500 bp upstream of the TSS and 500 bp downstream of the TES. *SCR* exons 1 and 2 are colored in black.

## Discussion

### S-locus sRNA precursors act through non-canonical gene silencing pathways

Our characterization of self-incompatibility transgenic lines of *A. thaliana* contributes additional evidence to the work of Durand et al. (19): collectively, these studies not only establish *A. thaliana* as a tractable system to study dominance interactions among *A. halleri S-*alleles, but also experimentally validate the role of two distinct sRNA precursor families in these interactions, even if the precise mechanisms underlying their action remain elusive. Our study reveals that some of the sRNAs generated in the processing of the *S*-locus sRNA precursors have a length of 24 nt and are loaded into AGO4, potentially indicating a role for RdDM. Nevertheless, *ago4 ago6*, *polIV* and *polV* mutations did not significantly affect the action of *A. halleri S-*locus sRNA precursors, suggesting that canonical RdDM is not the primary mechanism through which Ah20mirS3 and Ah04mir1887 mediate the reduction in *SCR01* transcript levels. While these results strongly suggest against DNA methylation as the basis of this transcriptional modulation, it would be interesting to directly measure DNA methylation at the *SCR01* locus using bisulfite sequencing or similar methods. However, such analyses remain challenging because dominance is established during early pollen development, specifically in tapetum cells, which are embedded within the anthers and difficult to isolate.

Considering their hairpin structure and their ability to also generate AGO1-loaded 21-nt sRNAs, it could also be envisioned that these precursors could act through PTGS. Still, the location of target sites in the promoter and intronic regions of *SCR01* poses a conundrum, since PTGS is reported to occur in the cytoplasm, using mature mRNA as a cleavage substrate (31). Notably, in animals, several lines of evidence point to miRNAs acting both in the cytoplasm and in the nucleus, with effects at the transcriptional and post-transcriptional levels (36–38). In plants, it has been observed that an inverted repeat targeting the intron of the soybean gene *FAD12-1A* triggers cleavage of its precursor mRNA (pre-mRNA) in the nucleus, which is accompanied by accumulation of siRNAs of the cleaved pre-mRNA (39). Additionally, all main miRNA processing components, including AGO1, are present in the nucleus (40–42). Thus, given the independent slicer activity of AGO1 (43), it could be hypothesized that this ARGONAUTE uses sRNAs to target the *SCR01* pre-mRNA in the nucleus, promoting its cleavage co-transcriptionally. Alternatively, the interaction between AGO1 and the *SCR* pre-mRNA could promote disassembly of the Poll II complex leading to the arrest of transcription, as has been suggested to occur in some miRNA genes upon salt stress (44). Even though we could not detect any siRNAs corresponding to the cleavage of the *SCR01* pre-mRNA in any of our sRNA sequencing experiments (data not shown), future experiments should investigate the possibility that the sRNA precursors at the *S-*locus could act through nuclear PTGS or through transcriptional arrest.

Our results indicating that Ah20mirS3 and Ah04mir1887 work through an RdDM-independent pathway contrast with previous observations in Brassica: in this latter system, the sRNA precursors *Smi* and *Smi2* are associated with deposition of DNA methylation in and around the *SCR* target site and a decrease in *SCR* expression (12, 20, 21). This could be a result from a direct incorporation of *Smi* and *Smi2* sRNAs into AGO4-clade proteins which would elicit *de novo* DNA methylation through the RdDM pathway, as has been observed for selected miRNAs and inverted repeats (22–26). Alternatively, as suggested by Finnegan et al. (45), it could depend on *Smi*-induced cleavage of an antisense transcript at the *SCR* locus, generating sRNAs that could subsequently be co-opted into the RdDM pathway. Even though empirical investigations are still necessary to test these hypotheses in Brassica, the cumulative evidence from these earlier studies and our stud suggests that different sRNA precursors within the Brassicaceae self-incompatibility system can recruit distinct effector pathways, that nevertheless converge into the same molecular phenotype: reduction of *SCR* transcript abundance. This would imply that these regulatory elements have an outstanding molecular flexibility, despite functioning within the specific evolutionary constraints of the self-incompatibility system. Thus, it cannot be excluded that different *A. halleri* sRNA precursors, as well as precursors from other self-incompatible species such as *A. lyrata* and *Capsella grandiflora* could use different gene regulatory pathways. Future investigations of different *S*-locus sRNA precursor families in these species, using heterologous expression systems as done in this study, or in the native context whenever genetic transformation is feasible, will hopefully help answer this outstanding question.

### The complex S-allele dominance network relies on the unique features of S-locus sRNA precursors

Durand et al. (19) previously showed that the *S-*locus sRNA precursors of *A. halleri* have high generalism, and suggested that dominant *S-*alleles in particular, rely on this feature to target multiple recessive alleles, rather than on carrying an increased number of sRNA precursors. Our study uncovers the molecular basis for this generalism, and shows that the apparent haphazardous processing of *S-* locus sRNA precursors is required to generate numerous functional sRNAs that target multiple alleles cooperatively: remarkably, all *SCR* alleles analyzed in this study are targeted by multiple sRNAs of a single sRNA precursor, and/or are targeted by distinct sRNA precursors. This contrasts with the less complex dominance hierarchy of Brassica, where fewer allelic interactions are present, and a single functional sRNA molecule per precursor has been suggested to mediate all dominance relationships (20, 21). It is important to note that while we investigated a total of 15 *S-*alleles in this study, at least 43 are known to exist in *A. halleri* (7), and > 80 in the closely related *Capsella grandiflora* (14). Therefore, in this study we most likely underestimated the fraction of functional sRNAs in each precursor, since those that are non-functional in the allelic interactions explored here could potentially have a role in interactions that have not yet been characterized.

Overall, the evidence suggests that collaborative targeting, either by multiple sRNAs of a single precursor, or by sRNAs from distinct sRNA precursors, is an essential feature of this system: firstly, it is essential for granting an *S*-allele its complete targeting spectrum against the many *S*-alleles segregating in the population; and secondly, the use of multiple sRNAs with distinct molecular features could trigger diverse effector pathways acting on the same target allele, which could perhaps underlie the robust reduction of recessive *SCR* transcripts observed in all allelic interactions studied thus far (this study and (10)).

Intriguingly, this resembles the collaborative non-self recognition model seen in Solanaceae, where each self-incompatibility allele contains multiple *SLF* genes. SLF proteins are essential for recognizing and neutralizing non-self S-RNases - female-derived ribonucleases that would otherwise inhibit pollen germination. In this system, the presence of multiple *SLF* genes within a single *S*-locus allele - each capable of recognizing a distinct S-RNase allele - together with the ability of some SLF proteins to recognize and detoxify multiple S-RNases (generalism), is crucial. Both these features ensure a broad recognition spectrum for each *S*-allele, preserving the complex network of inter-allelic interactions within Solanaceae (46). While SLF generalism in Solanaceae likely relies on the ability of each of these proteins to interact with multiple distinct S-RNases (47), sRNA precursor generalism in *A. halleri* relies on their unique molecular features, specifically their ability to produce numerous sRNAs with distinct targeting spectra.

### S-locus sRNA precursors provide a window into the miRNA evolution continuum

Although sharing some structural similarities with miRNAs, the features of *S*-locus sRNA precursors more closely resemble those of proto-miRNAs (48). Proto-miRNA loci are inverted repeats present in several plant genomes that have acquired transcriptional competence, forming long hairpins which can be diced by multiple Dicer-like proteins (DCLs), leading to a complex and heterogeneous population of sRNAs (48–52). There is little evidence that such a sRNA population could cause substantial effects in gene expression, at least through the canonical PTGS pathway; therefore, these structures have been mostly regarded as substrates for selection to act on, rather than having immediate biological significance (48–52). We hypothesize that the specific selective constraints of the *S*-locus dominance network have likely contributed to the persistence of proto-miRNA features over extended evolutionary times, and prevented their progression towards canonical miRNA features, as this would likely render them unable to effectively target multiple *S*-alleles. It would be interesting to test if apparently stabilized proto-miRNAs are also used in other biological systems where the expression of multiple genes/alleles needs to be coordinated by a single regulator and under similar evolutionary constraints as those acting on the *S-*locus. The mimicry of wing patterns in *Heliconius* butterflies would be an interesting model system to test this, since balancing selection has favored the appearance of numerous alleles of the supergene controlling wing pattern, and dominance relationships between them are prevalent (53, 54).

## Materials and Methods

### Plant material & growing conditions

Wild type (wt) and transgenic *Arabidopsis thaliana* plants used in this study were grown in a greenhouse, using standard growth conditions (16 h light/8 h dark; 110 μmol/s/m2; 21°C; 70% humidity). Seeds were germinated directly on soil, or in MS*-*medium (0.43% MS*-*salts, 0.8% Bacto agar, 0.19% MES hydrate, and 1% sucrose) when antibiotic selection was necessary for transgene selection. Because the self-incompatibility reaction cannot be faithfully reconstructed in the *A. thaliana* Col-0 ecotype (27–30) all wild type and mutant plants used in this study were of the C24 ecotype. *polIV* and *polV* correspond to the previously published and validated *rdm5 (nrpd1a)* and *rdm6 (nrpd1b*) mutant alleles (32). These two mutants were generated in the *ros1* mutant background, which was removed in our study by backcrossing to wt C24 plants. The *ago4 ago6* double mutant was generated in the C24 background using CRISPR/Cas9 **(see section *Generation and validation of CRISPR/Cas9 mutants* for more details)**.

To obtain the *S-*allele sequences of Ah65, Ah60, Ah03, Ah33 and Ah19, five *Arabidopsis halleri* individuals containing a combination of these alleles were collected from diverse natural European populations (55), and grown under standard greenhouse conditions in order to obtain plant material for DNA extraction and sequencing **(see the section *DNA extraction, Nanopore sequencing and assembly of S-alleles* for more details)**. Following Goubet et al. (15), we further obtained the S-allele sequence of Ah01, Ah02 and Ah25 by constructing and screening BAC libraries from fresh leaf samples, and fully sequencing the positive clones using PACBIO.

### Cloning

Several transgenic lines were used in this study: lines carrying SRK01, SCR01, and the sRNA precursor Ah20mirS3 were cloned as detailed previously (19).

The precursors Ah20mir1887 and Ah20mirS3 are in very close physical proximity within the Ah20 allele, and the previously published Ah20mirS3 line contains both sRNA precursors (19). While performing sRNA sequencing of AGO-IP experiments **(see section *ARGONAUTE immunoprecipitation and sRNA sequencing*)**, we noticed that some Ah20mir1887 sRNAs show homology to the 5’ region of *SCR01*. The base-pairing score of these predicted interactions varies between 12.5 and 18.5, depending on the sRNA **(see the section *sRNA database construction and target site inference for Ah04mir1887 and Ah20mirS3* for more details on these scores)**. To confidently isolate the effect of Ah20mirS3 in the dominance interaction of Ah20 over Ah01 (Interaction B), we introduced five point mutations in the SCR01^*mir1887^ sequence that disrupt the homology with the Ah20mir1887 sRNAs, rendering the Ah20mir1887 target site non-functional. The SCR01^*mir1887^ transgene was generated by performing site-directed mutagenesis on the already published SCR01 clone (19), using the primers detailed in **table S1**. The resulting amplicon was then recombined into the entry vector pDONR-Zeo, and subsequently recombined into the pB7m34GW destination vector using the 3-fragment Gateway Cloning System (Invitrogen), including 5’ and 3’ mock sequences, as detailed previously (19).

Ah04mir1887 was cloned by amplifying a region spanning this sRNAs precursor and containing 2 kb of upstream and downstream sequences from a BAC clone carrying allele Ah04, using the primers detailed in **table S1**. The resulting amplicon was then recombined into the entry vector pDONR-Zeo, and subsequently recombined into the pH7m34GW destination vector using the 3-fragment Gateway Cloning System (Invitrogen), including 5’ and 3’ mock sequences, as detailed previously (19).

Arabidopsis thaliana C24 plants were transformed using the floral dip method (56), and transgenic lines were selected using the appropriate antibiotics.

### Generation and validation of CRISPR/Cas9 mutants

Given the aforementioned limitation that the self-incompatibility reaction of *A. halleri* can only be reconstituted in the *A. thaliana* C24 ecotype (27–30), we created the *ago4 ago6* RdDM mutants in this specific ecotype using CRISPR/Cas9. The expression cassette of pCBC-DT1T2 (57) was amplified using two overlapping forward and reverse primers that contained two gene-specific sgRNA sequences (Fw1: 5’-ATATAT<u>GGTCTC</u>GATTGNNNNNNNNNNNNNNNNNNNGTT-3’; Fw2: 5’- TGNNNNNNNNNNNNNNNNNNNGTTTTAGAGCTAGAAATAGC-3’; Rv1: 5’- ATTATT<u>GGTCTC</u>GAAACNNNNNNNNNNNNNNNNNNNCAA-3’; Rv2: 5’-AACNNNNNNNNNNNNNNNNNNNCAATCTCTTAGTCGACTCTAC-3’, where N corresponds to the gene-specific gRNAs **(table S1)**, and underlined regions correspond to BsaI restriction sites). The amplicons obtained from this PCR reaction were gel-isolated and inserted into pHEE401E (58) using BsaI and T4 ligase (Thermo Fisher Scientific). The pHEE401E vectors containing gRNAs against *AGO4* and *AGO6* were transformed into the *Agrobacterium tumefaciens* strain GV3101. The floral-dip method (56) was then used to transform wt C24 *A. thaliana* plants.

To screen for T1 mutant plants, the regions targeted by the gRNAs were Sanger sequenced to identify *ago4 ago6* mutants with premature stop codons in both genes **(Fig. 1 S3)**. In the T2 generation, double homozygous mutants that did not carry the pHEE401E were selected by genotyping **(see table S1 for primers)**.

To confirm that *ago4 ago6* mutations impair DNA methylation, a Chop-PCR assay was performed (59). Briefly, this assay uses genomic DNA treated with the methylation-sensitive enzyme HaeIII as a template for PCR. Primers targeting regions known to have AGO4/AGO6-dependent DNA methylation, as well as control unmethylated regions are amplified **(see table S1 for primers)**. Using this method, we could determine that the *ago4 ago6* mutant generated in this study shows reduced DNA methylation levels, as has been previously published for other mutant alleles of these genes **(Fig. 1 S3 C)** (60).

### Pollen germination assays

All lines used for the reconstruction of the self-incompatibility reaction in *A. thaliana* were used in hemizygous state to mimic the abundance of sRNA precursors, and *SCR/SRK* in *A. halleri* individuals heterozygous at the *S-*locus. Genotyping of each transgenic line/mutant was done by classical PCR, or using the KASPar assay (61) **(table S1)**. Paternal plants hemizygous for *SCR01* and sRNA precursors were obtained by crossing *SCR01* to Ah04mir1887 homozygous lines or by crossing *SCR01^*mir^*^1887^ to Ah20mirS3 homozygous lines.

To assess the self-incompatibility reaction at the phenotypic level, pollen germination assays were performed on manually crossed plants. Maternal plants hemizygous for *SRK01* were emasculated one day before anthesis, and manually pollinated 24h later with paternal plants hemizygous for *SCR01, SCR01^*mir^*^1887^, *SCR01* Ah04mir1887 or *SCR01^*mir1887^* Ah20mirS3. Pollinated pistils were transferred to fixing solution 6h after pollination and stained with aniline blue, as described before (19). These pistils were then mounted on microscope slides and imaged using a Zeiss AX10 fluorescence microscope, where we counted the number of germinated pollen grains present in each pistil.

To evaluate the effect of RdDM mutants on the activity of *S-*locus sRNA precursors we crossed the RdDM mutants *ago4 ago6*, *polIV* and *polV* with the previously described paternal plants carrying *SCR01* and sRNA precursors (*SCR01, SCR01^*mir1887^*, *SCR01* Ah04mir1887 and *SCR01^*mir1887^* Ah20mirS3). We then obtained plants hemizygous for the *SCR01* and sRNA precursor transgenes and homozygous for the mutations of interest and used these as paternal plants in crosses with maternal hemizygous *SRK01* plants, as described above.

As a control, all maternal and paternal lines used in this study were crossed to wt C24. Germination and seed set for all these genotypes were comparable to that of wt x wt crosses (data not shown), confirming that the transgenes and mutations had no detectable effect on plant fertility.

### Phenotypic determination of the S-loci dominance hierarchy

The *S-*locus dominance hierarchy in **Figure 3** is based on a compilation of controlled reciprocal crosses between different *A. halleri* individuals (10, 19, 35, 62). In cases where interactions between alleles of two different dominance classes had not been tested phenotypically, dominance was inferred based on the phylogenetic class of each allele, since interclass dominance interactions are predictable (7, 16). In the case of intraclass relationships, such as for alleles Ah65, Ah60 and Ah33 (all belonging to class III), phenotypic dominance has not been determined yet, and thus the ordering of these alleles along the hierarchy remains arbitrary at this stage.

### RT-qPCR

To measure the abundance of *SCR01* transcripts, we isolated immature buds of plants hemizygous for *SCR01, SCR01^*mir1887^*, *SCR01* Ah04mir1887 or *SCR01^*mir1887^* Ah20mirS3. RNA was extracted using the NucleoSpin RNA Plus kit (Macherey-Nagel) and cDNA was synthesized with the RevertAid First Strand cDNA Synthesis Kit (Thermo Scientific). qPCR was performed using the iTaq Universal SYBR Green Supermix (BioRad) in a Lightcycler 480 instrument (Roche), with the primers found in **table S1**. *ACT8* was used as a reference gene. Transcript abundance was quantified using the Pfaffl method (63).

### ARGONAUTE immunoprecipitation and sRNA sequencing

AGO1 and AGO4 immunoprecipitation (IP) assays were performed on immature buds of *A. thalian*a plants carrying homozygous Ah04mir1887 or Ah20mirS3 transgenes, using a previously published protocol (64), with the following modifications: IPs were performed on 1.4 g of finely ground powder of frozen immature buds. The anti-AGO1 (AS09 527, Agrisera) and anti-AGO4 (AS09 617, Agrisera) antibodies were added at a 1/500 dilution for the incubation step and their immobilization was performed using Dynabeads Protein G (Invitrogen). The IP products were washed six times with PBS prior to sRNA extraction using the Trizol reagent (Ambion). In parallel, an aliquot of each input extract was used for total sRNA isolation using the Trizol LS reagent (Invitrogen), according to the supplier instructions. For each genotype three sRNA fractions were obtained: AGO1-IP, AGO4-IP and input.

These three sRNA fractions were subjected to an acrylamide gel-based size selection for sRNAs. These sRNAs were then used for TruSeq Small RNA (Illumina) library preparation and sequenced on a NextSeq 500 platform (Illumina), using 75 bp single-reads. Reads were trimmed, quality filtered, and size-selected using TrimGalore (65). We followed a two-step mapping procedure with ShortStack (allowing for one mismatch) (66), where we mapped sRNAs from each sample to the *A. thaliana* TAIR10 genome (masking the *S-*locus region), and to the sequence of the corresponding *A. halleri S-*allele (Ah04, for samples carrying Ah04mir1887 and Ah20 for samples carrying Ah20mirS3). Only reads that mapped exclusively to the *S-*locus were kept for further analysis. sRNA read counts were normalized to rpm to allow comparison of abundance levels across samples.

AGO loading was inferred based on the ratio of reads found in the AGO1-IP fraction and the AGO4- IP fraction: a sRNA was considered to be loaded in AGO1 or AGO4 when more than 50% of its total reads were derived from either the AGO1-IP fraction or the AGO4-IP fraction, respectively. A given sRNA was classified as having an unknown loading when no reads were found in both AGO-IP fractions, or when the amount of reads found in the input fraction was superior to that found in any of the two AGO-IP fractions.

### sRNA database construction and target site inference for Ah04mir1887 and Ah20mirS3

To identify sRNA target sites within *SCR* genomic regions, we first compiled a comprehensive database incorporating all sRNAs sequenced in the AGO-IP experiments described above, along with previously sequenced sRNAs (19). This database represents the complete set of sRNAs sequenced to date in our lab, and originating both from *A. thaliana* and *A. halleri* plants carrying Ah04mir1887 or Ah20mirS3. In parallel, a database containing *SCR* genomic regions of different *A. halleri S-*loci was compiled. This included alleles from all four dominance classes, and comprised previously published sequencing data **(see *Data Availability* section for a complete list of data sources)**. In addition to this, long read Nanopore sequences newly obtained from individuals carrying alleles Ah65, Ah60, Ah03, Ah33 and Ah19 were also added to this database **(see section *DNA extraction, Nanopore sequencing and assembly of S-alleles* for more details)**.

Using the databases of sRNA sequences and *SCR* alleles, we applied a previously published homology detection algorithm (19) to find target sites. Functional sRNAs are defined as those with a homology score of ≥ 18, since this score has been consistently shown to be associated with a decrease in *SCR* transcript abundance of the targeted allele (10, 19). The identified sRNA - *SCR* interactions were then represented in a heatmap **(Fig. 4)** using ComplexHeatmap (67), and in a circos plot **(Fig. 5)** using circlize (68).

### DNA extraction, Nanopore sequencing and assembly of S-alleles

To obtain the *S-*locus sequences of alleles Ah65, Ah60, Ah03, Ah33 and Ah19, 2 g of fresh leaves were collected from five different individuals carrying these alleles and flash-frozen. High molecular weight genomic DNA was extracted as described before (69). Long read sequences were obtained by Oxford Nanopore Technologies (ONT) flowcells v9. To polish the contigs obtained by long read sequencing Illumina data were also generated (with no step of PCR amplification to minimize sequencing bias).

For Nanopore library preparation, the smallest genomic DNA fragments were first eliminated using the Short Read Eliminator Kit (Pacific Biosciences). Libraries were then prepared according to the protocol 1D Native barcoding genomic DNA (with EXP-NBD104 and SQK-LSK109) provided by Oxford Nanopore Technologies. Depending on how many samples were pooled, 250 ng (pool of 9 samples) to 1 µg (pool of 4 samples) of genomic DNA fragments were repaired and end-prepped with the NEBNext FFPE DNA Repair Mix and the NEBNext Ultra II End Repair/dA-Tailing Module (New England Biolabs). Barcodes provided by ONT were then ligated using the Blunt/TA Ligase Master Mix (NEB). Barcoded fragments were purified with AMPure XP beads (Beckmann Coulter), then pooled and ONT adapters were added using the NEBNext Quick Ligation Module (NEB). After purification with AMPure XP beads (Beckmann Coulter), each library was mixed with the sequencing buffer (ONT) and the loading beads (ONT) and loaded on a PromethION R9.4.1 flow cell. In order to maintain the translocation speed, flow cells were refueled with 250 µl Flush Buffer when necessary. Reads were basecalled using Guppy version 3.2.10, 4.0.11, 5.0.12, 5.0.13, 5.0.16 or 5.1.12. The Nanopore long reads were not cleaned and raw reads were used for genome assembly.

For Illumina PCR-free library preparation, 1.5 μg of genomic DNA was sonicated to a 100–1500-bp size range using a Covaris E220 sonicator (Covaris). The fragments (1µg) were end-prepped, and Illumina adapters (NEXTFLEX Unique Dual Index Barcodes, Perkin Elmer) were added using the Kapa Hyper Prep Kit (Roche). The ligation products were purified twice with 1X AMPure XP beads (Beckman Coulter). The libraries were then quantified by qPCR using the KAPA Library Quantification Kit for Illumina Libraries (Roche), and their profiles were assessed on an Agilent Bioanalyzer (Agilent Technologies). The libraries were sequenced on an Illumina NovaSeq 6000 instrument (Illumina) using 150 base-length read chemistry in a paired-end mode.

Nanopore sequencing data were assembled using Necat (70) with a genome size of 240 Mbp and the remaining parameters left as default. Contigs produced by Necat were polished one time using Racon (71) with Nanopore reads, then one time with Medaka (https://github.com/nanoporetech/medaka, model r941_prom_hac_g507) and Nanopore reads, and two times with Hapo-G v1.3.4 (72) and Illumina short reads. However, the cumulative sizes of our long-read assemblies were larger than the estimated 240 Mb, suggesting that the assembly size was currently inflated by the presence of allelic duplications. Thus, HaploMerger2 (73) was run (Batch A twice to remove major misjoin and one Batch B) to generate a haploid version of each assembly.

To detect the *S-*locus region in the generated assemblies, blast searches were performed using the genes flanking the *S-*locus (*ARK3* and *U-box* (15)) as queries. This allowed us to precisely narrow down the genomic regions corresponding to the full Ah65, Ah60, Ah03, Ah33 and Ah19 *S-*alleles. We annotated *SRK* using blast searches, and following Goubet et al. (15), we used Fgenesh+ (74) to predict the genomic sequence of *SCR* based on a database of SCR protein sequences from already known S-alleles.

### Determination of the SCR01 transcription start site (TSS)

To identify the TSS of *SCR01* in *A. halleri*, total RNA was extracted from unopened buds of three individuals plants from accession Zapa12 (Central Europe), Bara01 and Firi03 (Romania), genotyped as homozygous for the Ah01 allele. A library was directly prepared using the NEBNext Ultra II Directional RNA Library Prep for Illumina Kit. Libraries were sequenced with an Illumina NovaSeq 6000 instrument (Illumina) using a paired-end 151 bp read chemistry. Short Illumina reads were bioinformatically processed to remove adapters and primer sequences, only reads ≥ 30 nucleotides were retained. These filtering steps were done using in-house-designed software based on the FastX package (75). Finally, read pairs that mapped to ribosomal sequences were filtered using SortMeRNA as previously described (76). Processed reads were aligned with STAR (77) using –alignIntronMax 3000 and –alignMatesGapMac 10000 parameters to avoid very long splicing assignment errors due to a mammalian-based design of the STAR algorithm. The vertical coverage at each base pair for the three replicates was computed using samtools depth (78) and then plotted in R after feature scaling and smoothing the curves using mean coverage values in 20 bp sliding windows.

### Data availability

The AGO-IP and sRNA sequencing data is deposited in the Gene Expression Omnibus database, under the SubSeries reference GSE249507.

The Illumina and Oxford Nanopore sequencing data of the five *A. halleri* individuals carrying alleles Ah65, Ah60, Ah03, Ah33 and Ah19 are available in the European Nucleotide Archive, under the project number PRJEB70880.

The RNA-seq data used to infer the *SCR01* transcription start site are available at ENA-EBI under the following accession numbers ERR14765081, ERR14765082 and ERR14765083.

The *S*-locus sequences used in this study are a combination of previously published data (15, 19) and data generated in this study. All allele sequences can be found in GenBank, under the following references: Ah01 - this study, PP396894; Ah02 - this study, PP396895; Ah03 - this study, PP537897; Ah04 - KJ461484; Ah10 - KM592810 and KM592817; Ah12 - KJ772373 and KJ772377; Ah13 - KJ461479 and KJ461483; Ah19 - this study, PP537898; Ah20 - FO203486; Ah25 - this study, PP396896; Ah28 - KJ461478; Ah29 - KM592803; Ah33 - this study, PP537899; Ah60 - this study, PP537900; Ah65 - this study, PP537901.

## Acknowledgments

This work was supported by a European Molecular Biology Organization (EMBO) Postdoctoral Fellowship attributed to RAB, by the European Research Council (NOVEL project, grant #648321) and by the French National Research Agency (TE-MoMa project, grant ANR-18-CE02-0020-01). This work was performed using the infrastructure and technical support of the Plateforme Serre, Cultures et Terrains Expérimentaux - Université de Lille for the greenhouse/field facilities. Research at the Lagrange laboratory was supported by the Centre National de la Recherche Scientifique (CNRS) and Université de Perpignan Via Domitia (UPVD). This study is set within the framework of the “Laboratoires d’Excellences (LABEX)” TULIP (ANR-10-LABX-41) and the “Ecole Universitaire de Recherche” (EUR) TULIP-GS (ANR-18-EURE-0019). Research at the Krämer laboratory was supported by the Deutsche Forschungsgemeinschaft DFG in SPP 1529 (ADAPTOMICS, Kr1967/10-1,-2), and by ERC-AdG LEAP-EXTREME (788380). We thank Hervé Vaucheret, Vincent Colot, Claudia Köhler and Nicolas Butel for helpful discussions, and Hervé Vaucheret for sharing seeds.

## Author Contributions

Designed research – RAB, ED, MM, JA-F, MD, SS, VK, IF-L, MB, J-MA, SL, UK, TL, XV, VC; Performed research – RAB, ED, MM, JA-F, MD, SS, VK, EL, CC, CB, MB, A-CH, CP, NF, IF-L; Contributed new reagents or analytic tools – WM, SV, IF-L, J-MA, UK, TL, XV, VC; Analyzed data – RAB, ED, MM, JA-F, MD, SS, VK, EL, MB, J-MA, SL, XV, VC; Wrote the paper – RAB, VC. All authors reviewed and edited the manuscript.

---

**Figure 1 S1.**
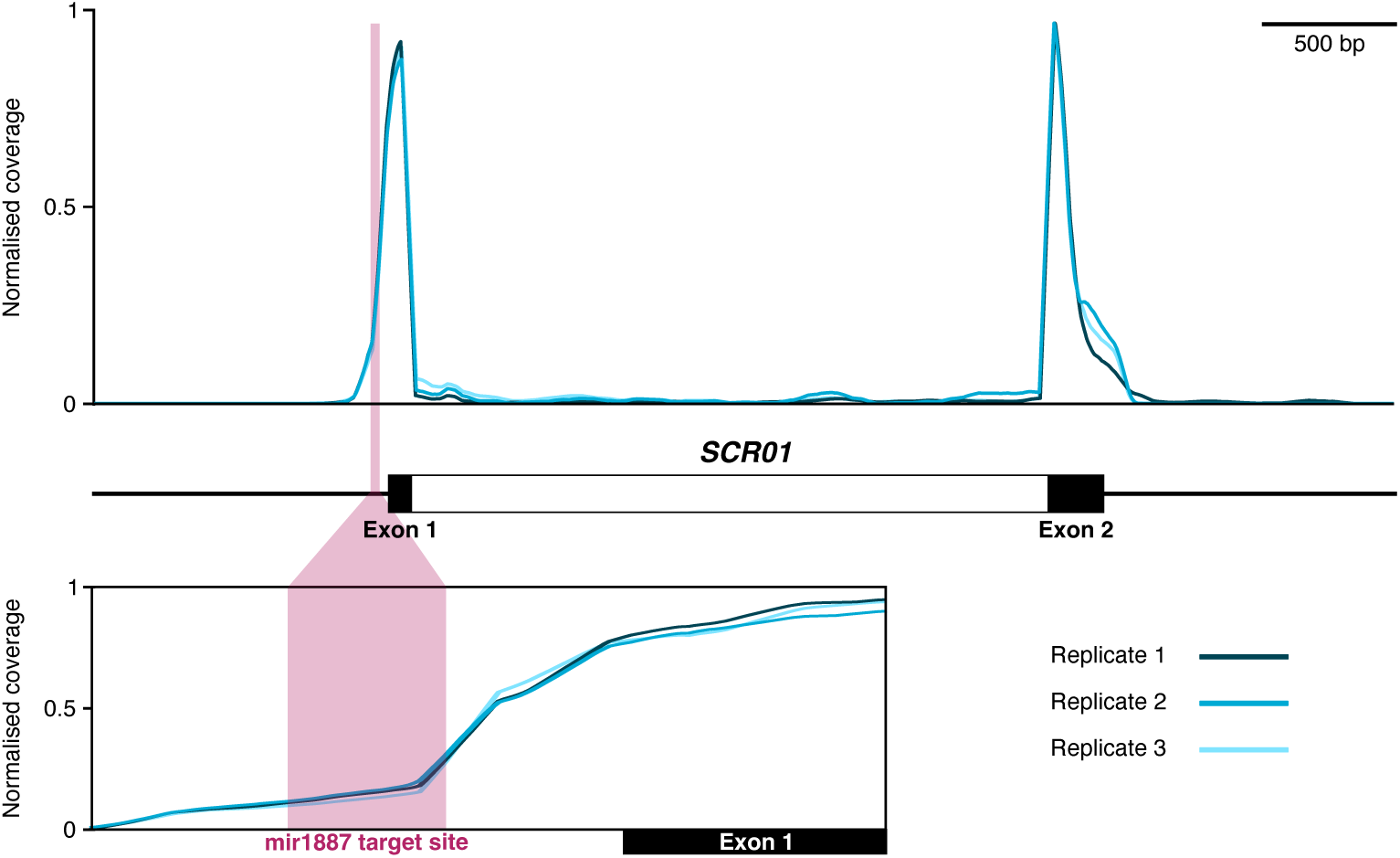
SCR TSS in relation to Ah04mir1887 target site. Normalised RNA-seq coverage along the *SCR01* sequence from three biological replicates of homozygous *Ah01* allele plants. Exons and introns within *SCR01* gene are represented by black and white regions, respectively. The *mir1887* target site is highlighted in light red and the bottom panel provides a magnified view of this site, including the flanking exon 1. The three replicates are depicted as curves in different shades of blue. Raw vertical coverage distributions (number of aligned reads at each position) are normalized to the maximum value for each replicate. The curves are smoothed using a 20 bp sliding window, advancing by 1 bp per step.

**Figure 1 S2.**
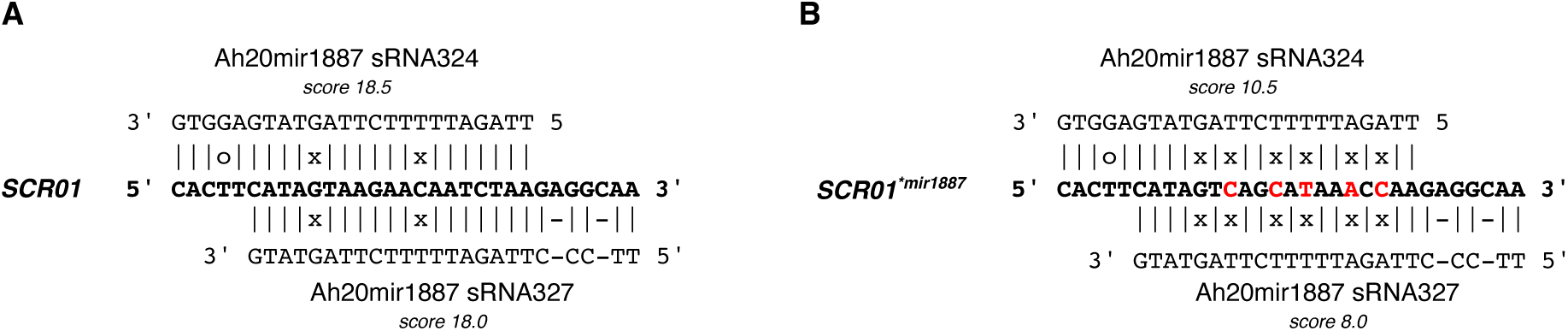
*SCR01^*mir1887^* transgenic line. **(A)** Alignment between the native mir1887 target site on SCR01 and Ah20mir1887 sRNAs with a targeting score ≥ 18. **(B)** Alignment between the same sRNAs and the mutated mir1887 target site of the transgenic line SCR01*mir1887. Note that the mutations decrease the homology score below 18, suggesting that these sRNAs are not able to target the mutated site. Mutated sites are in red.

**Figure 1 S3.**
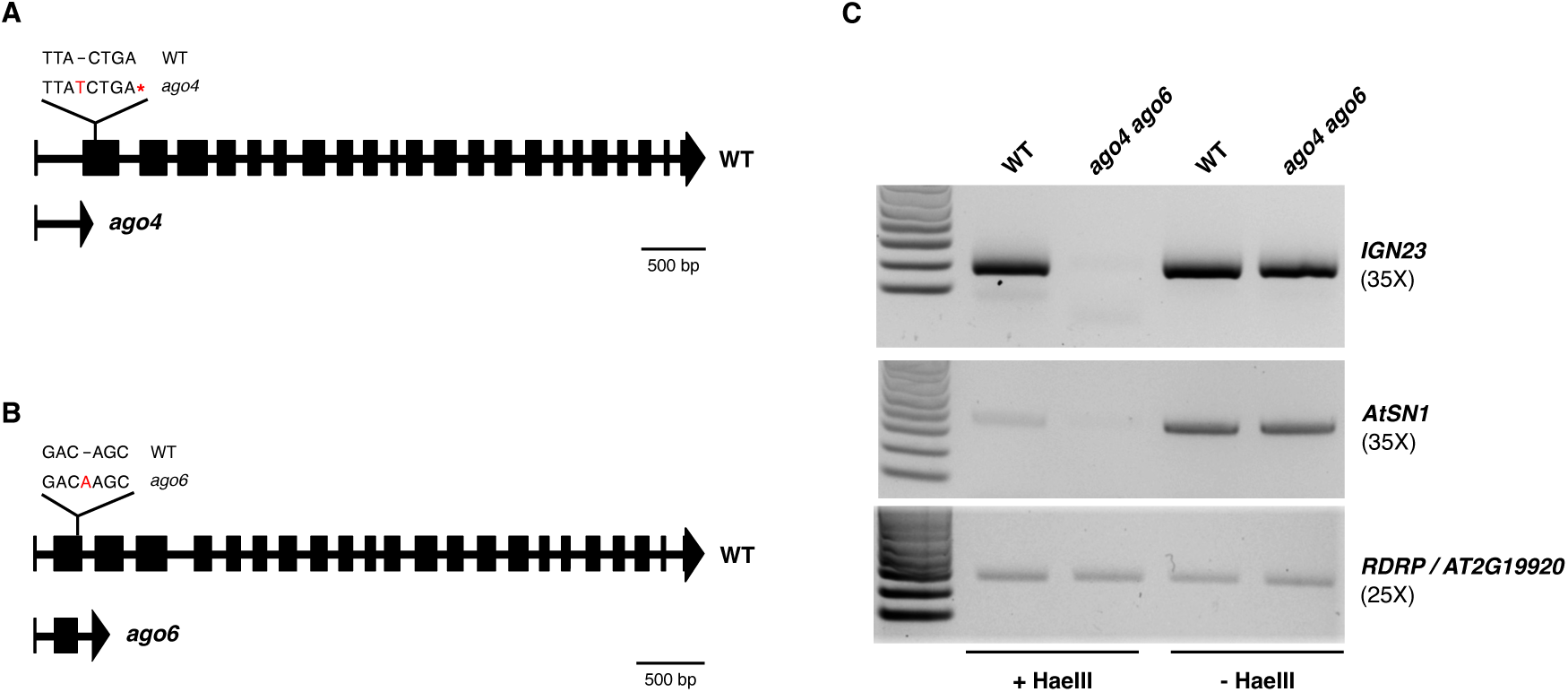
Validation of the RdDM CRISPR/Cas9 mutants generated in this study. **(A-B)** Schematic representation of CRISPR/Cas9 induced mutations in the *AGO4* and *AGO6* genes. Sequence changes are shown in red, premature stop codons are represented by the asterisk. **(C)** Chop-PCR assay on wt C24 plants and *ago4 ago6* double mutant plants. *IGN23* and *AtSN1* correspond to RdDM-dependent methylated regions, while *RDRP* corresponds to a non-methylated region.

**Figure 1 S4.**
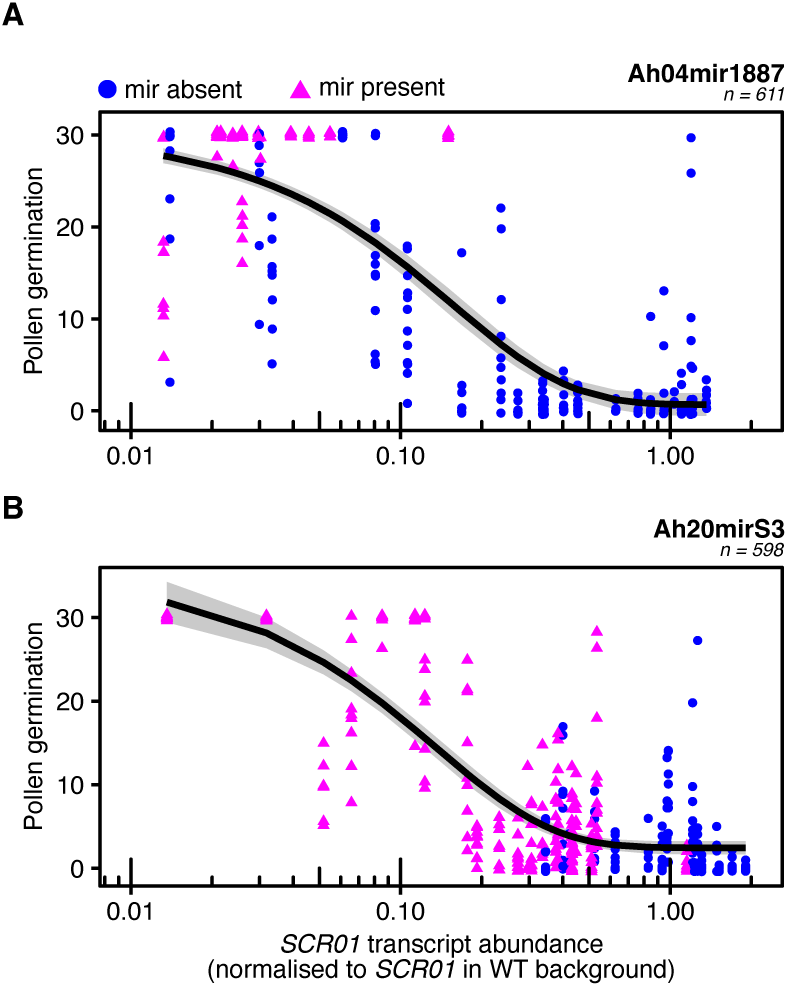
A compatible pollen germination phenotype requires a low *SCR01* transcript abundance. Non-linear regression analysis modeling the relationship between *SCR01* transcript abundance levels and pollen germination. For both Ah04mir1887 **(A)** and Ah20mirS3 **(B)** sRNA precursors a compatible pollen germination phenotype (∼20-30 pollen grains) requires a 10-fold decrease in *SCR01* transcript abundance (compared to the level observed in the wt). Analysis was performed using the phenotypic and expression data presented in **Fig. 1 C-D**. The black line represents the predicted model and the gray area corresponds to the 95% confidence interval.

**Figure 2 S1.**
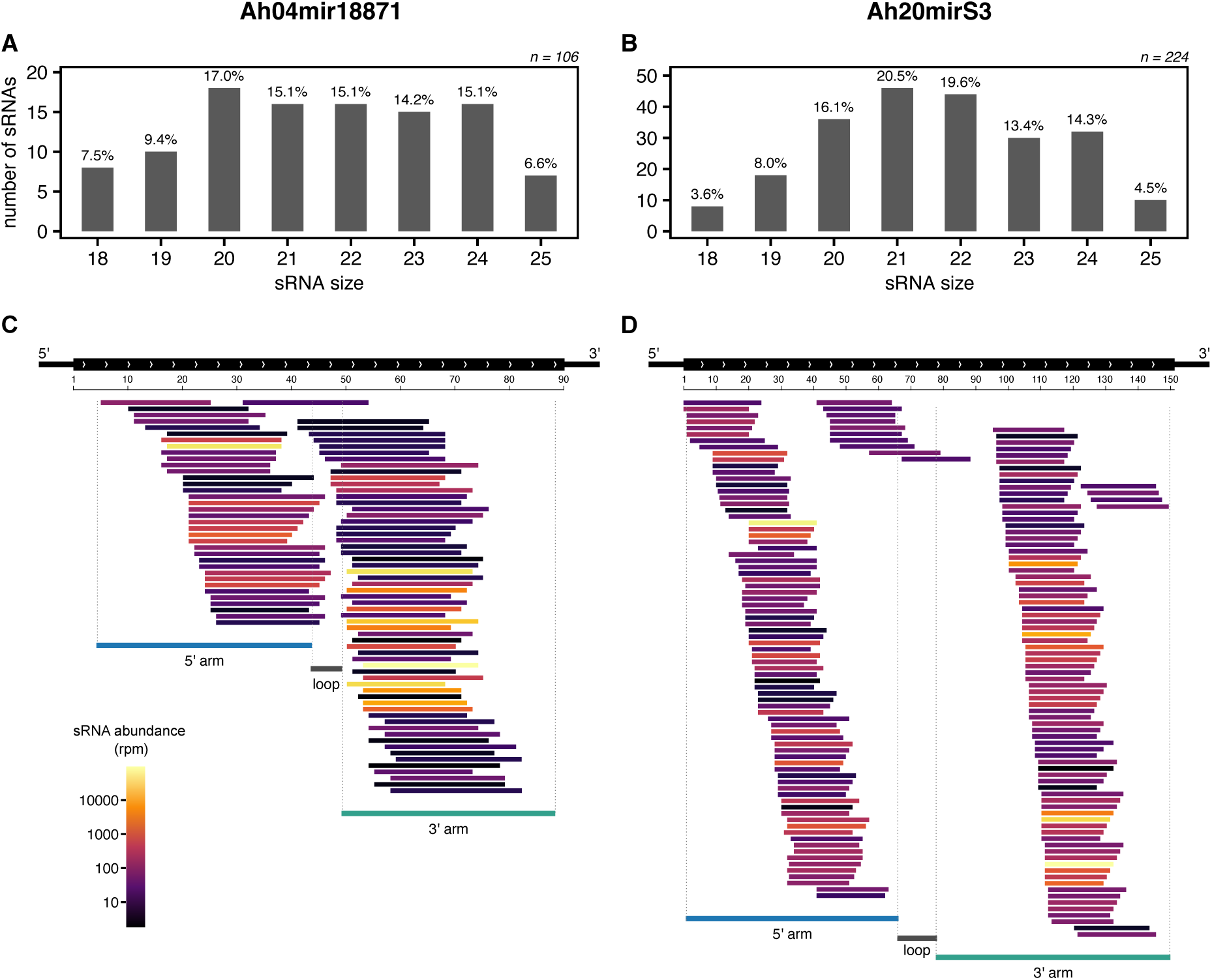
sRNAs produced from Ah04mir1887 and Ah20mirS3 do not show a specific size bias but vary in their abundance levels. Size distribution of sRNAs derived from Ah04mir1887 **(A)** and Ah20mirS3 **(B)**. **(C-D)** Genomic representation of the Ah04mir1887 and Ah20mirS3 loci, and their respective sRNAs. sRNAs are colored according to their abundance level.

**Figure 2 S2.**
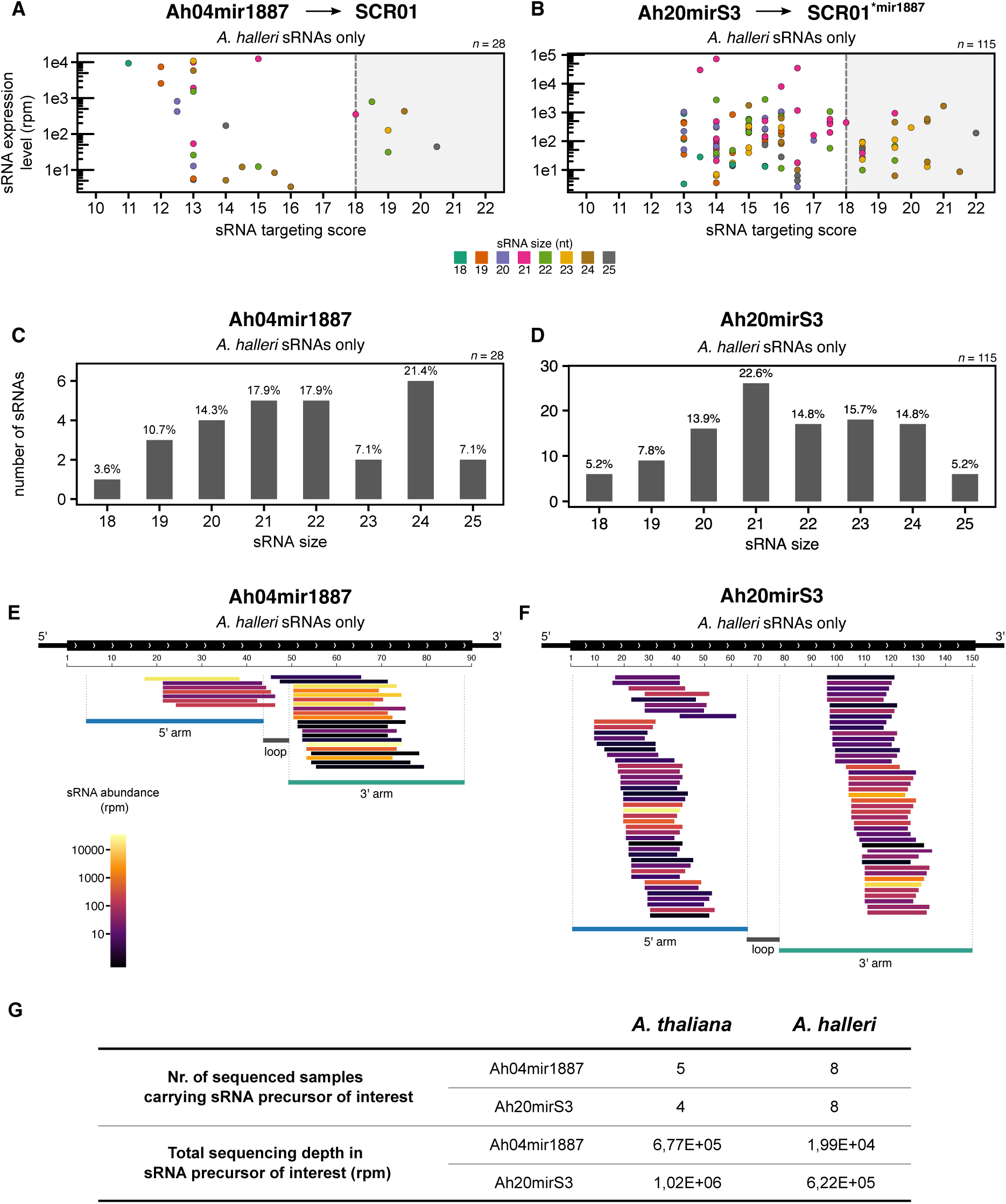
Molecular features of *A. halleri*-specific sRNAs produced by Ah04mir1887 and Ah20mirS3. **(A-B)** Expression of Ah04mir1887 **(A)** and Ah20mirS3 **(B)** *A. halleri* sRNAs as a function of their targeting score against *SCR01.* The dot color corresponds to the size of the sRNA. The grey box highlights sRNAs predicted to induce a reduction in *SCR01* transcript abundance (score ≥ 18). *n* corresponds to the total number of unique sRNAs identified in *A. halleri*. **(C-D)** Size distribution of *A. halleri* sRNAs derived from Ah04mir1887 **(C)** and Ah20mirS3 **(D)**. **(E-F)** Genomic representation of the Ah04mir1887 **(E)** and Ah20mirS3 **(F)** loci, and the respective *A. halleri* sRNAs. sRNAs are colored according to their abundance level. **(G)** Summary of sRNA sequencing efforts in *A. thaliana* transgenic lines and *A. halleri* lines carrying Ah04mir1887 and Ah20mirS3.

**Figure 2 S3.**
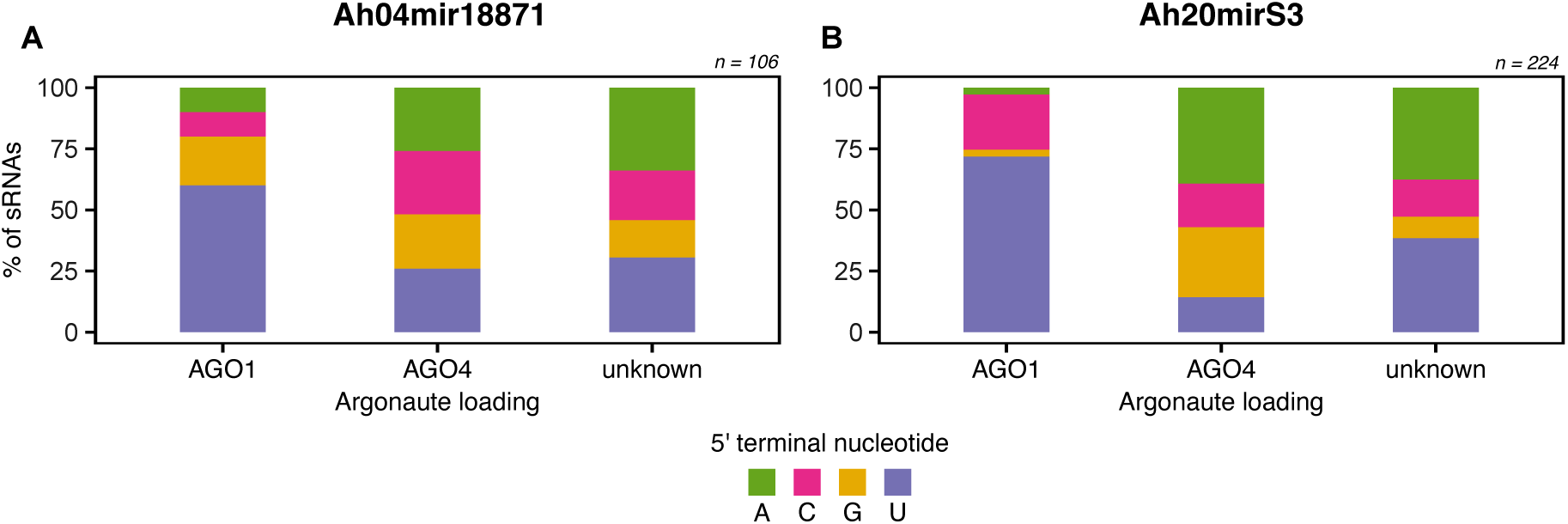
5’ terminal nucleotide frequencies in Ah04mir1887 (A) and Ah20mirS3 (B) sRNAs, related to their inferred AGO loading pattern.

